# Deep-ultraviolet microscopy reveals biomolecular spatiotemporal intracellular dynamics

**DOI:** 10.64898/2026.08.02.742310

**Authors:** Viswanath Gorti, Mingxuan Si, Nathan Taylor, Marcus Cicerone, Francisco E. Robles

## Abstract

Intracellular dynamics span a broad range of time scales and biomolecular processes, offering insights into cell health, functional state, phenotype, and response to external perturbations. Several label-free optical imaging approaches have been used to capture intracellular dynamics but are limited by spatiotemporal resolution and biomolecular specificity required to distinguish unique subcellular and metabolic processes. In this work, we demonstrate deep-ultraviolet (UV) microscopy as a powerful, label-free, high-resolution approach for quantifying multiscale intracellular dynamics with biomolecular specificity. By leveraging power spectral analysis and phasor analysis, we capture multiscale intracellular dynamics and analyze their UV wavelength-dependent behavior predicated by the absorption of different endogenous biomolecules. We apply this technique to prostate epithelial cell lines of increasing malignancy and reveal quantitative differences in dynamic intracellular activity that correlate with increased metabolic and organelle activity between phenotypes. Furthermore, we elucidate the molecular identities of structures and activity measured via UV dynamics with broad-band coherent anti-Stokes Raman scattering spectroscopy and fluorescence microscopy. We identify lipid-specific structures, and mitochondrial-specific dynamics, among other biomolecular-specific dynamic behaviors. Together, this study demonstrates deep-UV microscopy as a powerful imaging platform for probing spatial and temporally variant intracellular dynamics with biomolecular specificity, with broad implications for cell phenotyping, tissue pathology, and studying new dynamic subcellular processes.

## Introduction

Intracellular dynamics, including processes such as organelle motility, membrane fluctuation, vesicle trafficking, and nuclear dynamics, are critical for cell function. These dynamic phenomena occur across a broad range of spatial and temporal scales, from macro-scale cell growth and motility over seconds to nucleic acid and protein fluctuations on the order of milliseconds (1–3). Variations in intracellular dynamics are associated with cellular metabolic activity and functional state, making them key indicators for assessing cell health, subtype, and response to external perturbations. Quantifying these dynamics can provide insight into fundamental biological processes and metabolic profiles, enable new applications for non-destructive cell phenotyping.

To fully capture intracellular dynamics, an imaging technique must be able to operate at a wide range of temporal scales, be non-perturbative to the sample (i.e., label-free), and offer targeted biomolecular specificity to distinguish intracellular activity. Several optical imaging techniques have been developed to visualize and quantify intracellular dynamics. Fluorescence-based methods, such as confocal fluorescence microscopy and fluorescence correlation spectroscopy, have been demonstrated to collect intracellular dynamics with high biomolecular specificity (2–4). However, these rely on fluorescent labels, which can perturb native cell dynamics, introduce artifacts, and are often limited by photobleaching and cytotoxic effects.

To combat these limitations, label-free imaging modalities have been applied for dynamic live cell imaging. Quantitative phase imaging (QPI) non-destructively measures refractive index fluctuations in tissue to track changes in cell morphology and dry mass over time. Recent work with QPI has demonstrated the ability to capture intracellular fluctuations corresponding with metabolic activity at low frame rates (5–8). Dynamic full-field optical coherence tomography (D-FFOCT) has also been used to monitor intracellular motion within organoids and dense tissue samples (9–11). Furthermore, Raman-based techniques provide biochemical specificity without labeling, allowing for molecular fingerprinting of live cells (12–17).

However, these label-free techniques have key limitations for longitudinal measurement of intracellular dynamics. QPI and OCT lack biomolecular specificity, relying on refractive index and backscattering to generate contrast within cells, inhibiting the specificity of the dynamic information. While chemical imaging systems do offer biomolecular specificity, they often suffer from long acquisition times, low throughput, or poor spatial resolution, limiting the breadth of intracellular dynamics that can be captured.

Deep-ultraviolet (UV) microscopy enables label-free, and fast imaging of live cells with targeted biomolecular specificity, offering a promising approach for capturing multiscale intracellular dynamics. Deep-UV microscopy leverages strong absorption of endogenous biomolecules in the 200-300 nm spectral range, generating high-resolution images with intrinsic molecular contrast (18–21). Notably, nucleic acids exhibit a peak absorbance around 255 nm, while proteins absorb strongly at 220 and 280 nm due to peptides and aromatic amino acids, respectively. Lipids, collagen, and other macromolecules also demonstrate absorption in this region, making deep-UV microscopy a versatile technique for capturing bio-molecular features within cells (21). The use of UV light for biological imaging was first demonstrated over one hundred years ago, but was limited due to inefficient UV imaging hardware, leading to concerns over UV-induced photodamage during live cell imaging studies. However, recent advances in UV optics - including illumination sources, and high quantum efficiency detectors – enable fast label-free imaging at biologically relevant wavelengths. As a result, deep-UV microscopy has been demonstrated for several hematology and histopathology applications (22–33). More recently, deep-UV imaging has been shown as a non-destructive and fast approach for high-throughput immune cell characterization, (34). We have also recently demonstrated that with proper dose management, deep-UV microscopy can be applied for longitudinal cell studies without causing detectable photodamage (33).

In this work, we advance deep-UV microscopy by imaging and quantifying intracellular dynamics over a broader set of imaging conditions and cell types. Using multispectral UV illumination and high-speed imaging, we capture intracellular activity at both slow (0.5-5 Hz) and fast (5-50 Hz) temporal scales, leveraging frequency-domain analysis to quantify measured activity. We apply this technique to several prostate epithelial cell lines of varying malignancy to demonstrate wave-length- and frequency-dependent differences in measured dynamics, capable of distinguishing unique cell types that are morphologically similar. Furthermore, we validate the molecular sources of deep-UV dynamic contrast via broadband coherent anti-Stokes Raman scattering (BCARS) spectral imaging. Together, this study highlights the potential of deep-UV microscopy for quantifying multiscale intracellular dynamics with high specificity, throughput, and simplicity, with broad implications for phenotyping cell type and functional state.

## Results

### Deep-UV microcopy enables visualization of intracellular dynamics

To demonstrate the capability of deep-UV microscopy for visualizing intracellular activity in live cells, we used a previously developed benchtop UV microscopy setup capable of high-speed, label-free imaging at biologically relevant UV wavelengths (22). The system comprises a laser-driven plasma broadband light source, with narrowband UV bandpass filters, a sample stage containing adherent cells cultured in petri dishes, and a UV objective (40X, 0.5NA) delivering light to a UV-sensitive camera (Fig. 1A). This configuration enables rapid acquisition frame rates up to 1 kHz with ~300 nm spatial resolution, sufficient to resolve subcellular features (further described in Methods). While this study used a benchtop UV microscope with research-grade hardware, we note that similar imaging capability (i.e., high-resolution, multispectral deep-UV microscopy) can be achieved with previously demonstrated low-cost, LED-based, compact UV microscopy systems (26, 34).

**Fig. 1:**
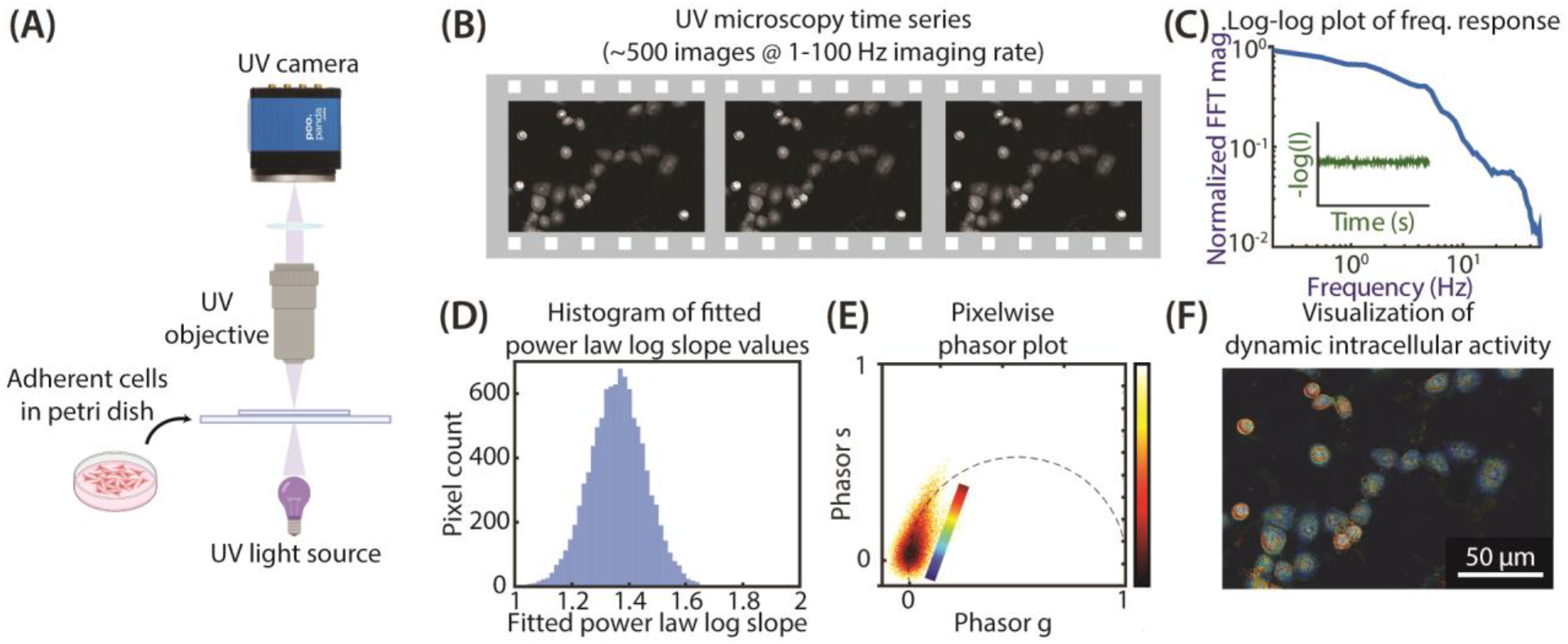
Deep-UV microscopy enables quantitative analysis of intracellular dynamics. (A) Schematic of imaging protocol with adherent cells imaged in a deep-ultraviolet (UV) microscopy setup. (B) Representative images of RWPE-1 cells under 255 nm illumination at 100 Hz imaging frame rate during a deep-UV microscopy time series. (C) Log-log plot showing the normalized frequency response of pixel values (representing attenuation) in a segmented cell (inset shows a sample time profile of pixelwise attenuation). (D) Histogram of pixelwise fitted power law log slope values for a single cell. (E) Phasor plot of extracted pixelwise phasor g and s values for a single cell. (F) Phasor-colorized pseudo colorized image of RWPE-1 cells visualizing intracellular dynamic activity under 255 nm illumination at 100 Hz imaging frame rate.

Representative frames from a time-lapse image stack captured with 255 nm illumination at 10 Hz imaging rate are shown in Fig. 1B, in which intensity fluctuations over time indicate intracellular activity. To quantify these dynamics, we extracted pixelwise intensity time series from segmented cell regions (Fig. 1C), which demonstrated heterogenous temporal fluctuations. To visualize frequency-based activity, we performed a Fourier-based power spectral analysis on each pixel’s temporal signal creating linear and log-log representations of intracellular dynamics (Fig. 1C). Recent works, using QPI and an optical tweezers-based approach, have proposed that intracellular dynamics follow a power law decay model in the frequency domain with no dominant oscillatory behaviors, suggesting predominantly stochastic Brownian and advective intracellular dynamics (5, 35). The dynamic signals are expected to follow a frequency dependance of *I*(*f*) ∝ *f*^−*β*^, where *β* ~ 1 indicates diffuse (Brownian) motion whereas a *β* ~ 2 indicates advective cellular motion (5, 35).

To quantify intracellular dynamics, we performed two complementary analyses. First, we fit each pixelwise frequency response to a power law decay curve, with the fitted log slope serving as an indicator of the dynamic activity parameter *β*. The fitting range and corresponding slope calculation are further detailed in the methods section. A histogram of these log-slope values reveals a distribution approximately between one and two, as expected (Fig. 1D). We then applied phasor analysis to the pixelwise frequency response, a technique commonly used for decomposing spectral signals (36–38), generating two terms denoted *g* and *s* which correspond to the real and imaginary components of the signal’s (*I*(*f*)) Fourier Transform at a given period. Using these terms, we created a phasor plot – a 2D histogram of *g* and *s* values that provide a graphical representation of the dynamics (Fig. 1E). Points further from the origin (0,0) in phasor space indicate higher temporal fluctuation and thus greater measured activity. Dynamics that map further from the origin in phasor space also exhibit a *β* closer to 2, indicating more advective cellular motion. Points closer to the origin in phasor space exhibit a lower *β* indicating more Brownian motion. Finally, to facilitate interpretation of these dynamics, we pseudo colorized cells with a colormap corresponding with where the dynamics at each spatial pixel map to in phasor space (Fig. 1F). The resulting map reveals heterogeneous dynamic regions within cells, corresponding with intracellular dynamics, including organelle motion, protein trafficking, and lipid vesicle transport. The information not only enables detection of functional intracellular activity but also provides insights into the cells’ biomolecular and phenotypic composition.

### Measured intracellular dynamics vary with imaging wavelength and sampling frequency

To investigate the wavelength- and frequency-dependence of intracellular dynamics, we imaged live prostate epithelial cells using illumination from three deep-UV wavelengths – 239, 255, and 280 nm – which approximately correspond with absorption peaks of lipids, nucleic acids, and proteins, respectively. For each wavelength, we acquired image stacks at two distinct frame rates (10 and 100 Hz), corresponding with effective dynamic frequency ranges of approximately 0.5-5 Hz and 5-50 Hz, respectively (see Methods). This enabled interrogation of both slow and fast intracellular activity with varying functional contrast.

Figure 2 shows results from RWPE-1 cells, which are non-tumorigenic epithelial cells derived from adult human prostate. This cell line was immortalized with HPV-18 and mimics normal prostate epithelium, expressing no tumorigenicity or pathologies (39). The top row of Fig. 2 shows UV attenuation images (-log of the transmission intensity) across the three wavelengths used in this study, revealing consistent cell and nuclear morphology with some differences in subcellular and cytoplasmic contrast.

**Fig. 2:**
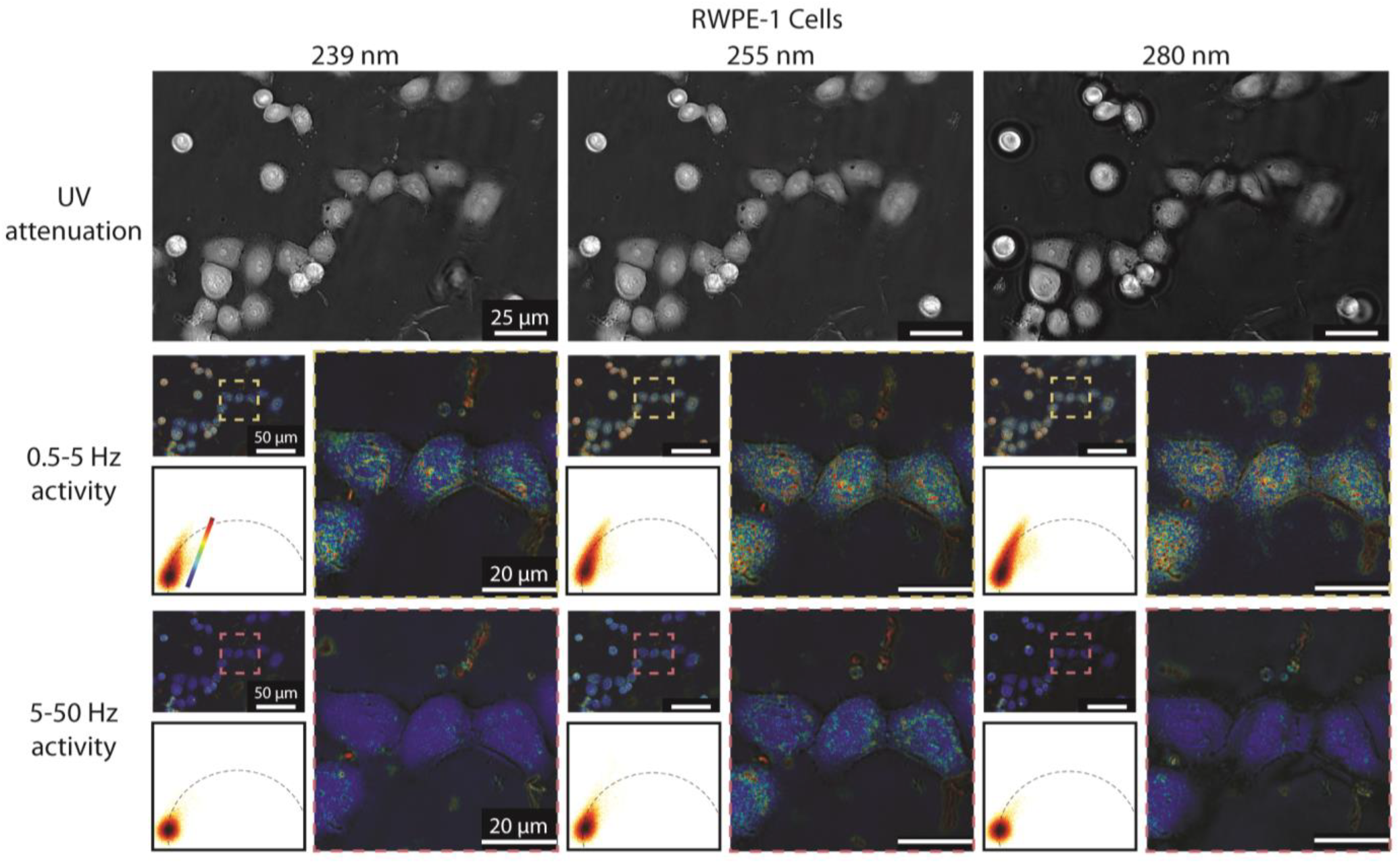
Measured intracellular dynamics in RWPE-1 cells vary with UV illumination wavelength and activity frequency range. UV attenuation images of adhered RWPE-1 cells with 239 nm (left), 255 nm (center) and 280 nm (right) illumination. Phasor-colorized images, phasor plots, and cropped insets shown for activity ranges corresponding with low (middle row) and high (bottom row) frequency activity ranges. The 0.5-5 Hz activity range corresponds with 10 Hz imaging frame rate, while the 5-50 Hz activity range corresponds with 100 Hz imaging frame rate.

Phasor analysis revealed differneces in measured dynamic activity that varied with both wavelength and sampling frequency. In the 0.5-5 Hz activity range (10 Hz imaging frame rate; Fig. 2, middle row), we observed moderate dynamic activity in RWPE-1 cells, particularly under 255 and 280 nm illumination. The phasor plots show the spread of measured activity within segmented cell regions, while the pseudocolorized images highlight fluctuations in the cytoplasmic regions. Notably, cell and nuclear membrane regions demonstrated some dynamic activity but a lower amplitude of activity was present within the nucleus itself with the exception of the nucleoli. The activity measured with 239 nm illumination was generally weaker and more diffuse, consistent with broader but less specific absorption properties of biomolecules at this band.

In contrast, the 5-50 Hz activity range (100 Hz imaging frame rate; Fig. 2, bottom row) exhibited reduced relative intracellular activity across all wavelengths. Note that all pseudocolorizations were created with the same colorscale range. Interestingly, we observed more tightly packed clusters in phasor space closer to the origin, which indicates overall reduced high-frequency dynamics. Nevertheless, notable activity was present under 255 nm illumination, localized again to cytoplasmic compartments, but was substantially lower relative to the 0.5-5 Hz activity range. These dynamics also appear to be localized to small “hotspots”. Given the nucleic acid and protein source of contrast at this wavelength, we posit that these structures—not readily appreciable in static images but clearly revealed via dynamic analysis—are mitochondria in the cell cytoplasm (this is further discussed below). Frequency response plots corresponding with phasor-colorized activity (e.g., low activity pixels colored blue and high activity pixels colored red) corroborate these differences in measured activity (Supplemental Fig. S1). In the 0-5Hz activity range, pixels within each activity bin show significant difference in average frequency response, suggesting that some intracellular regions are more dynamic throughout this frequency range. In the 5-50Hz activity range, only pixels colorized red, corresponding with high dynamic activity, demonstrate a significantly different frequency response. These correspond with the aforementioned “hotspots”, revealed via dynamic analysis, which are the only intracellular structures with high frequency activity.

To investigate whether measured intracellular dynamics vary in cells with different activity profiles, we repeated the same analysis with WPE1-NB26 cells, a more aggressive cell line derived from RWPE-1 cells treated with a direct-acting carcinogen (*N*-methyl-*N*-nitrosoure, commonly known as MNU), that exhibit higher tumorigenicity and invasive characteristics (39). We hypothesized that the aggressive and less differentiated nature of these cells, resulting in faster doubling time and greater metabolic activity, would provide differing dynamic signal (40–42). Attenuation images of these cells (Fig. 3, top row) demonstrate broad morphological differences including increased proliferation and elongated cell structure, as expected with the malignant cell line. Dynamic activity in the 0.5-5 Hz activity range was consistently elevated relative to RWPE-1 cells, with increased measured activity in cytoplasmic regions across all three wavelengths. This increase suggests more active molecular dynamics and transport activity across all three wavelengths. Interestingly, the 5-50 Hz dynamics in WPE1-NB26 cells also showed a smaller but still apparent increase in activity under 255 and 280 nm illumination, with phasor plots demonstrated expanded clusters compared to RWPE-1 cells. The presence of more distinguishable high-frequency activity in WPE1-NB26 cells suggests that measured fast dynamic activity may correlated with malignancy-associated cell functions.

**Fig. 3:**
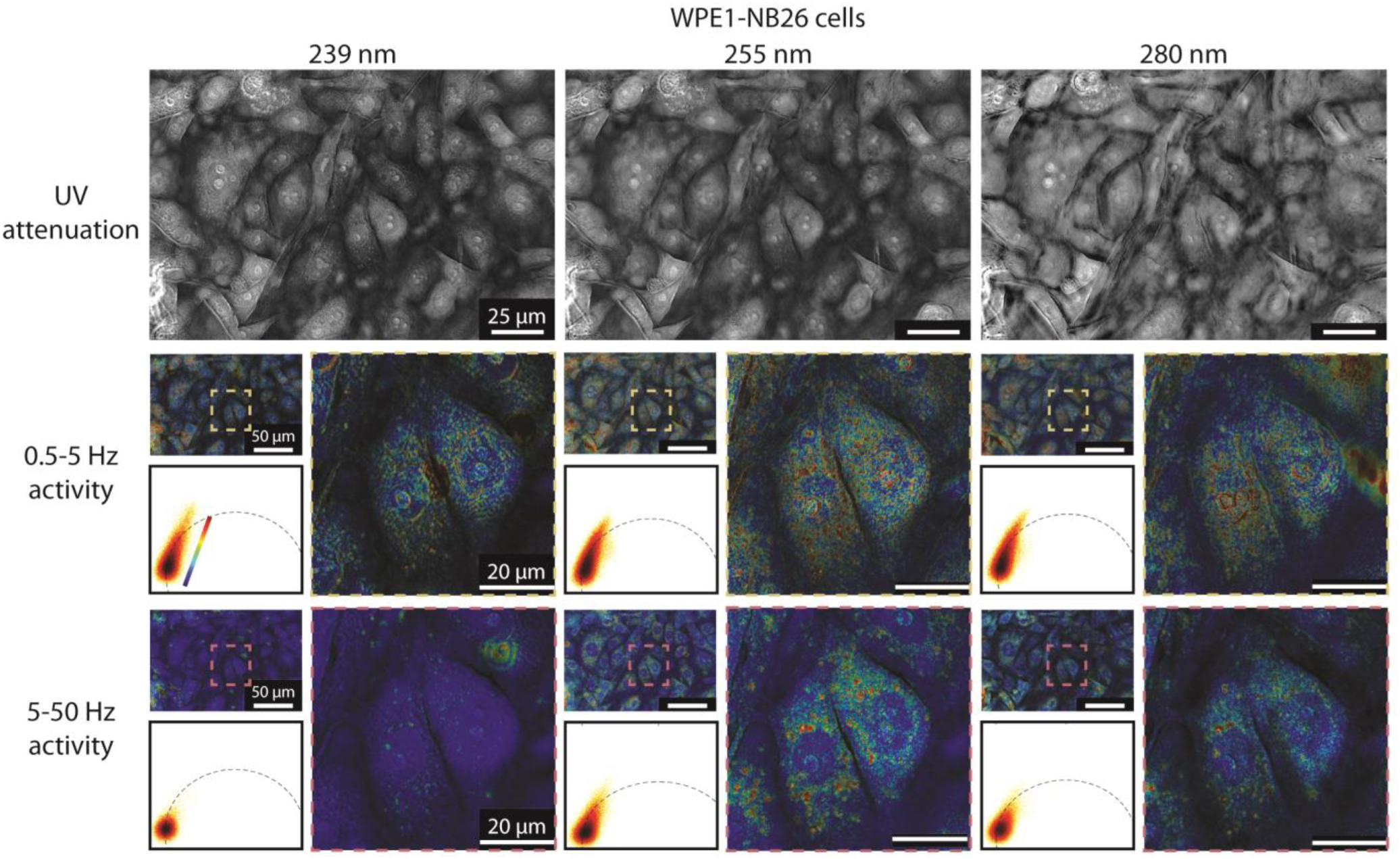
Measured intracellular dynamics in WPE1-NB26 cells vary with UV illumination wavelength and activity frequency range. UV attenuation images of adhered WPE1-NB26 cells with 239 nm (left), 255 nm (center) and 280 nm (right) illumination. Phasor-colorized images, phasor plots, and cropped insets shown for activity ranges corresponding with low (middle row) and high (bottom row) frequency activity ranges. The 0.5-5 Hz activity range corresponds with 10 Hz imaging frame rate, while the 5-50 Hz activity range corresponds with 100 Hz imaging frame rate.

More importantly, the 5-50 Hz pseudo-colorized images acquired with 255 nm and 280 nm illumination reveal a high density of well-defined activity “hotspots”, again these are also seen in non malignant RWPE-1 cells but with a lower density. Again, we posit that these structures correspond to mitochondria. Mitochondria are dynamic organelles that undergo continuous dynamic processes, including fission, transport, and nucleoid remodeling – all of which occur on fast time scales within this detectable frequency range (43). Previous live cell imaging studies have shown that mitochondrial nucleoid dynamics, including attachment and detachment, occur several times per second across a cell (44). Furthermore, mitochondrial fusion and fission cycles, including mitochondrial transport, all involve subcellular processes and interactions occurring at sub-second rates (despite these cycles lasting several seconds in total) (44–46). These processes are indeed related to cell health and physiology, and could vary with cell phenotype (47), including with cancer where the more fragmented mitochondria in increasignly more malignant cancer cells exhibit increased motility and hence dynamics(48).

The 5-50 Hz pseudo-colorized image with 239 illumination also shows interesting results. The static attenuation image, shows bright (more attenuating) spots, which likely correspond to lipids (not mitochondria) as lipid absorption in the deep-UV region is high in this range, with absorption peaks varying with lipid type and oxidation state (49–51). However, the static image alone can be ambigious. Conversely, analysis of the same dynamics shows little to no dynamic activity. This rules out that the static structures seen in the absroption images are mitochondria, and supports the notion that the dynamics observed using 255 nm illumination do not arrise from lipids, which are expected to have much slower dynamics (in the order of seconds to minutes) (1–3).

To better visualize the spatial organization and heterogeneity the dynamic “hotspot”structures, Fig. 4 presents a larger-FOV view of high-frequency intracellular dynamics in WPE1-NB26 cells under 255 nm illumination. This visualization reveals that dynamic hotspots are distributed throughout the cytoplasm showing perinuclear enrichment (localization around the nuclear membrane). This behavior is consistent with organelle-associated cellular activity near the nuclear envelope. Furthermore, there is some cell-to-cell heterogeneity in the number and location of these hotspots, which varies with cell health and age. Indeed, these hotspots are difficult to visualize in static UV attenuation images (Fig. 3) alone because of the small size and low concentration of the chromophore, both of which provide subtle contrast in static images. However, by monitoring changes over time, constrast to these dynamic structures is significantly amplified, underscoring the value of multiscale dynamic UV imaging for revealing subcellular activity.

**Fig. 4:**
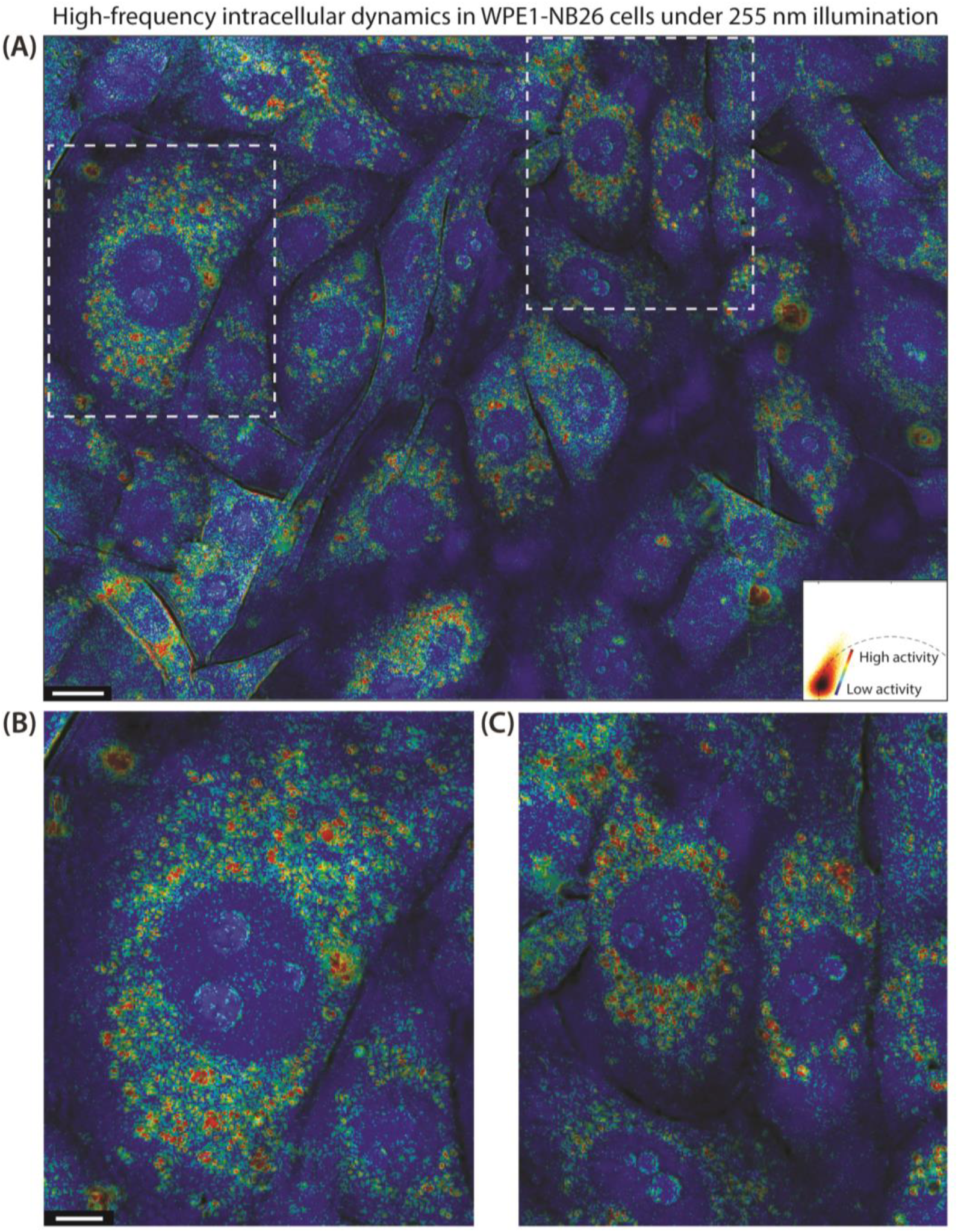
High-frequency intracellular dynamics in WPE1-NB26 cells under 255 nm illumination. (A) Large field-of-view phasor-colorized map of high-frequency (5-50 Hz) intracellular dynamics in adhered WPE1-NB26 cells under 255 nm UV illumination at 100 Hz imaging frame rate. Color represents relative dynamic activity magnitude with warmer (red) colors indicating higher magnitude of measured activity. (B-C) Zoomed-in views of representative regions of (A) highlighting individual cells and notable puncta demonstrating higher activity. Phasor color mapping is identical across all panels. Scale bars: 10 μm (A); 5 μm (B, C).

Furthermore, we collected image stacks at 1 Hz imaging rate to investigate very low-frequency (<0.5 Hz) dynamics that have been reported in other works (5). However, under these conditions, the observed signal was dominated by motion artifcats (e.g., floating debris, general whole cell movement and growth), and did not yield significant intracellular activity. These data are included in Supplemental Fig. S2, but were excluded from downstream analyses.

### BCARS imaging and fluorescence microscopy validate biomolecular contrast in deep-UV microscopy images

To further validate the contrast observed in deep-UV microscopy images and confirm the biomolecular composition of intracellular structures of interest, we performed broadband coherent anti-Stokes Raman scattering (BCARS) imaging of RWPE-1 cells. BCARS provides highly specific information of the chemical composition of cells with high resolution (12–14). Notably, we aimed to identify the composition of small, round, highly absorbing structures observed in a subset of adherent cells, suspected to be lipid droplets, which are most prominent in the 239 nm illumination attenuation images. Fig. 5A and 5E (and insets in Fig. 5B and 5F) show comparisons of deep-UV attenuation at 239 nm illumination and BCARS signal at 2930 cm^-1^ for the same field of view containing multiple cells. Morphologically, the two modalities reveal similar contrast and intracellular morphology, including nuclei, nucleoli, and cytoplasm, highlighting the biomolecular specificity of UV microscopy without complex optical setups or cell fixation.

**Fig. 5:**
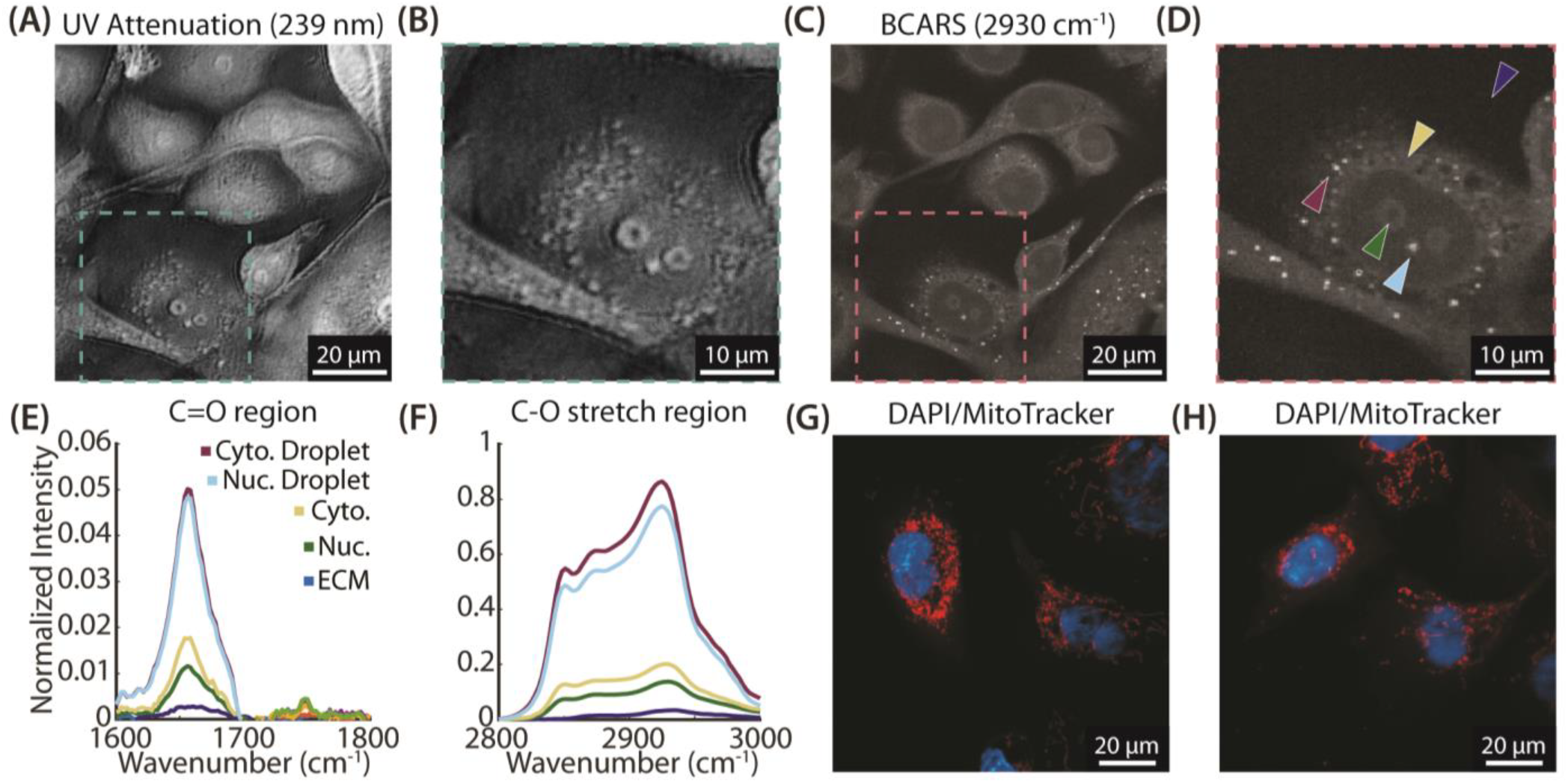
BCARS imaging validates biomolecular contrast from subcellular structures in deep-UV microscopy images. (A) Deep-UV attenuation image at 239 nm showing RWPE-1 cells. (B) Inset of (A) highlighting a well-adhered cell with highly absorbing subcellular droplets. (C) BCARS intensity image at 2930 cm^-1^ of the same RWPE-1 cells shown in (A) after fixation. (D) Inset of (C). Arrows indicate pixels corresponding with the extracellular matrix (ECM, dark blue), nucleus (green), cytoplasm (gold), a nuclear droplet (light blue), and cytoplasmic droplet (maroon). (E) BCARS spectra from the points in (D) in the ester C=O region (1600-1800 cm^-1^), showing strong peaks for cytoplasmic and nuclear droplets. Cytoplasm, nucleus, and ECM show reduced intensity. (F) BCARS spectra from the C-H stretch region (2800-3000 cm^-1^) of the same pixel locations. (G, H) Fluorescence images of RWPE-1 cells stained with DAPI (blue, nuclei) and MitoTracker Red (mitochondria) showing typical mitochondrial network morphology.

We further interrogated these structures by extracting and plotting pixelwise BCARS spectra at notable regions, including the extracellular matrix (ECM), cytoplasm, nucleus, and other structures of interest in the nucleolus and cytoplasm. The selected points are indicated with arrows in the BCARS inset (Fig. 5F). Plots of BCARS spectra at these points corresponding with the ester C=O carbonyl region and C-O stretch region are provided in Fig. 5C-D (full spectra are provided in Supplemental Fig. S3). In both regions, the extracellular region exhibited very low signal, as expected. Pixels corresponding with the nucleus and the cytoplasm exhibited slightly higher signal, consistent with some lipid and protein content. Notably, the droplets demonstrated a very strong signal in both spectral ranges, corresponding with high lipid content. Specifically, high signal in the carbonyl region (~1600-1800 cm^-1^), including increased signal at the ester carbonyl peak (1748 cm^-1^), suggests an abundance of triglycerides (TGs) and cholesteryl esters (CEs). Lipid droplets, vesicles, and endosomes are dense in TGs and CEs, while other organelles (e.g., mitochondria) can be rich in phospholipids but are relatively sparse in these lipid subtypes (60). Moreover, these cytoplasmic and nuclear lipid bodies are not present in all cells and often appear in cells with altered morphologies. Accumulation of neutral lipids (e.g., TGs and CEs) is an adaptive protection mechanism and can be a response to cell stress, during which toxic lipid intermediates are neutralized to prevent lipotoxicity (61).

To further validate that these structures were not mitochondria, we performed fluorescence imaging with DAPI and MitoTracker Red to stain nuclei and mitochondria, respectively. The resulting images reveal the typical branching mitochondrial network distinct from the distinct droplets observed in the UV and BCARS images (Fig. 5G-H). Thus, we conclude that the absorbing structures observed in the 239 nm attenuation images are in fact lipid-rich subcellular structures (such as lipid droplets, vesicles, or endosomes), corroborating recent studies on cellular activity using other imaging modalities (6).

These results are also consistent with the dynamic images taken at 100 Hz using 239 nm illumination, which do not show significant dynamic activity. This further supports that the 5-50 Hz dynamics observed with 255 and 280 nm illumination (which are dominated by absorption from nucleic acid and proteins, respectively) do not originate from lipid dynamics. Instead, the observed structure is more consistent with the spatial distribution of mitochondria—as seen in MitoTracker Red fluorescence images (Fig. 5H)—which contain both nucleic acid and proteins. The more heterogeneous dynamic contrast at lower frequency dynamics (0.5-5 Hz) also provides some phenotypic differentiation, but it is likely that many other dynamic processes are being captured at these lower frequencies, whereas the high frequency dynamics appear to be more specific to mitochondria.

These results demonstrate that deep-UV microscopy is capable of visualizing intracellular biomolecular heterogeneity without the need for chemical labels, fixation, complex optical setups, or lengthy acquisition times. We note that BCARS is a very powerful and highly specific biochemical imaging modality, but deep-UV microscopy can provide important advantages in certain use cases given its simplicity, low cost, and speed. Further multi-modal studies with UV microscopy and BCARS (or other imaging methods) can enable identification and quantitative measurement of other unique subcellular organelles and molecules, offering a powerful, label-free approach assessment of intracellular functional and molecular activity.

### Quantitative static and dynamic features reveal differences between cell phenotypes

After establishing the capability of deep-UV microscopy for detecting spatiotemporally variant intracellular dynamics, we investigated whether the functional information is associated with differences in cell phenotypes. Specifically, we aimed to assess if cellular and intracellular structures and dynamics differ between RWPE-1 cells and malignant derivatives/phenotypes. To do this, we extracted over 1000 quantitative features per cell, including (1) static morphological features (e.g., area, circularity); (2) static textural features (e.g., entropy, homogeneity) calculated from UV attenuation images at the three key wavelengths; (3) cellwise dynamic features corresponding with each illumination type and imaging frame rate (e.g., mean cellwise phasor g/s value and pixelwise power law log slope coefficient); and (4) textural features using pixelwise dynamic maps (e.g., entropy of a pixelwise phasor s map). These features have been extensively detailed in prior works for cell and tissue characterization (52–58).

We hypothesized that morphology alone would be sufficient to distinguish between the extreme phenotypes of benign RWPE-1 cells and aggressive WPE1-NB26 cancer cells; thus, we also analyzed an intermediate phenotype, WPE1-NB11 cells, derived from the same RWPE-1 lineage but with lower tumorigenicity than WPE1-NB26 cells (39). This enabled us to explore what dynamic features contribute to phenotype separation, and thus isolate the specific, underlying functional processes contributing to this differentiation. To identify the significant features, we performed chi-squared feature ranking across all features and classes and selected eight features for analysis. The resulting clustergram revealed clear separation of cells by phenotype with several features (including static and morphological features) contributing to the separation (Fig. 6A).

**Fig. 6:**
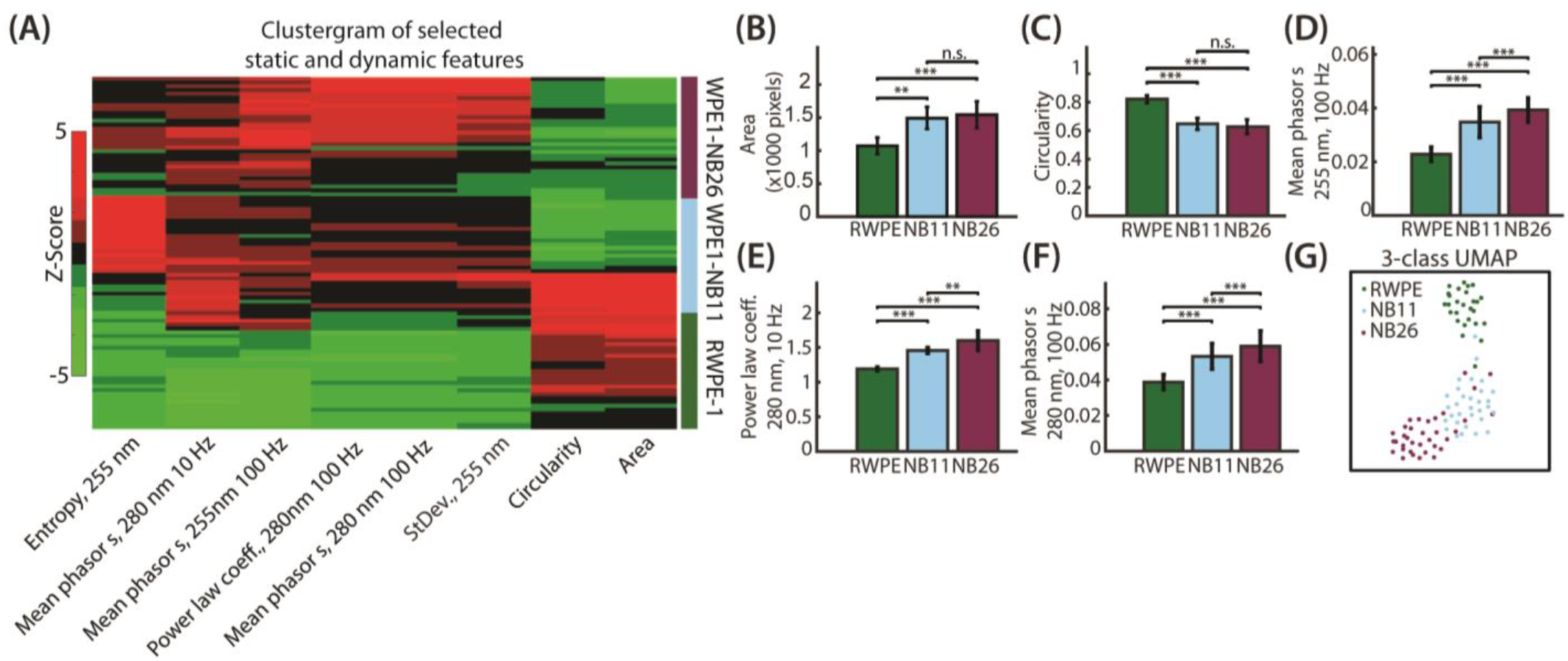
Quantitative static and dynamic features distinguish prostate epithelial cell phenotypes. (A) A 3 class clustergram of the top eight features (identified via chi-squared feature ranking) for classification of prostate epithelial cell phenotypes. Each row represents a single cell and columns are z-score values for each feature. The colored bar on the right indicates cell class: RWPE-1 (green), WPE1-NB11 (blue), and WPE1-NB26 (purple). (B-C) Bar plots of mean cell area (B) and circularity (C) per class show significant differences between RWPE-1 cells and both derived malignant cell lines, but not between WPE1-NB11 and WPE1-NB26 cells (mean +/-s.d.) (D-F) Bar plots of dynamic features including mean phasor s value at 255 nm, 100 Hz imaging rate (D), power law log slope coefficient at 280 nm, 10 Hz imaging rate (E), and mean phasor s value at 280 nm, 100 Hz imaging rate (F) show increased activity from RWPE-1 cells to subsequent derived cell lines, with all pairwise comparisons significant (mean +/-s.d.). (G) UMAP visualization of all cells using the eight features shown in (A) reveals clear phenotypic separation between RWPE-1, WPE1-NB11, and WPE1-NB26 cells. ** Student’s *t*-test p-value < 0.01, *** Student’s *t*-test p-value < 0.001. N = 101 cells.

We then explored the behavior of individual features in the different cell phenotypes. Morphological differences, including cell size and circularity, were significantly different between benign RWPE-1 cells and the cancerous phenotypes, but showed no significant difference between the less aggressive WPE1-NB11 and more aggressive WPE1-NB26 cells (Fig. 6B-C). This is consistent with known characteristics of cancer cell progression, where cells become larger and more elongated due to cytoskeletal remodeling and invasive behavior (40, 59). However, these structural changes are not significantly different between the phenotypes with different levels of malignancy (i.e., metastatic potential).

In contrast, dynamic features revealed significant differences across all three cell lines (Fig. 6D-F). Specifically, mean cellwise phasor s values at 255 nm/100 Hz, cellwise power law log slope coefficient at 280 nm/10 Hz, and mean phasor s values at 280 nm/100 Hz all increase with malignancy (i.e., from RWPE-1 to WPE1-NB11 to WPE1-NB26 cells). These features reflect biomolecular dynamics at nucleic acid and protein dominant absorption wavelengths, and at both slow (0.5-5 Hz) and fast (5-50 Hz) temporal frequencies, corroborating the trends seen between Fig. 2 and Fig. 3 above. The increased measured dynamic activity in the more aggressive phenotypes for all these features is due to a combination of multiple factors, including increased metabolic activity (resulting in more cytoplasmic remodeling, organelle transport), dynamic changes in fragmented mitochondria, and other subcellular phenomena characteristic of metastatic cancer cells (5, 6, 40, 41).

Indeed, deep-UV microscopy provides a wide range of both biomolecular-specific structural and dynamic/functional information that can be used to phenotype cells in an accessible configuration. To assess how well these features can distinguish cell phenotypes when combined, we visualized all cells in a 3-class UMAP using the features from the clustergram in Fig. 6A (Fig. 6G). The resulting 2D representation showed clear separation among RWPE-1, WPE1-NB11, and WPE1-NB26 cells. Notably, RWPE-1 cells formed a distinct cluster far from both malignant phenotypes, with the more metastatic cell line being furthest away from the benign group. The two malignant cell phenotypes, WPE1-NB11 and WPE1-NB26 cells, also mapped to unique locations but in closer proximity to one another. This pattern results from the static and dynamic differences discussed above and closely reflects the tumorigenicity characteristics of these cells. Ultimately, this visualization highlights the capability of deep-UV microscopy to extract multiscale, biologically relevant features from live cells that differentiate unique cell phenotypes by capturing biomolecular structure and activity.

## Discussion

In this work, we establish deep-UV microscopy as a powerful and accessible tool for functional and molecular imaging with the ability to quantify intracellular dynamics and cell phenotypes. Moreover, by imaging at key deep-UV biomolecular absorption wavelengths and capturing dynamic fluctuations across a wide temporal time-scales, we show that wavelength and temporal-specific activity can be used to distinguish between benign, intermediant, and highly aggressive prostate cancer cell lines. Importantly, the ability to resolve unique intracellular dynamics suggests that dynamic deep-UV microscopy can be applied for characterization of a wide range of cellular conditions, beyond cell malignancy.

Unlike other label-free techniques, such as QPI, which rely on broad refractive index differences within biological samples, deep-UV microscopy leverages intrinsic biomolecular absorption to enable contrast from nucleic acids, proteins, lipids, and other molecules. This specificity enables localized measurement of subcellular structures and processes occurring within them. Specifically, we show that intracellular dynamics at 255 and 280 illumination reveal differences in intracellular dynamics that correlated with malignant cell behavior, such as increase metabolic activity, organelle movement and altered protein and nucleic acid dynamics. These differences are quantified with higher phasor values and steeper power law decay slopes, consistent with prior studies investigating metabolic activity within cancer cell phenotypes (5, 35). Notably, we demonstrate differences in measured dynamic activity between closely related malignant cell lines (e.g., WPE1-NB11 and WPE1-NB26) that do not exhibit significant morphological differences.

Furthermore, our findings highlight that dynamic intracellular activity is also differentiable by frequency range. By imaging at 10 and 100 Hz (corresponding with activity ranges of approximately 0.5-5 Hz and 5-50 Hz, respectively), we observed frequency-dependent differences in activity between cell lines. (A long acquisition (100s) acquired at high frame rates (100 Hz) can capture both the “low” and “high” frequency dynamics but this is much more memory intensive as it requires 10,000 acquisitions, see Supplemental Fig. S4.) While RWPE-1 cells exhibited minimal activity in the high-frequency band, derived malignant cells demonstrated increased activity in the high frequency range, suggesting that aggressive cell phenotypes may also have more rapid subcellular fluctuations, reflecting increased metabolic flux and activity. We also note, that the frequency dynamics generally exhbit a power law behavior, without any resonant peaks in the measured frequency range, which can be well characterized by phasor analysis, as well as a power law fit within a frequency range (see Supplemental Fig. S4). Future work, can extend the power law analysis to modle more complex behavior across the measured spectrum.

Importantly, we find that the dynamic image analysis, particularly at high frame rates, reveals intracellular structures (e.g., mitochondria), that could not be easily distinguished in static UV attenuation images. While intracellular dynamics were heterogenous at lower frame rates (0.5 – 5 Hz), higher frame rate dynamics revealed small hotspots within cells, consistent with fast mitochondrial dynamics. These dynamics are not significantly present in data collected with 239 nm illumination, corresponding primarily with lipid absorption, but appear strongly with UV illumination correspond with nucleic acid and protein absorption – two compounds that mitochondria are rich in. Moreover, this suggests that the source of phenotyping contrast between cell lines of varying malignancy is in part due to mitochondrial dynamics, which corroborates literature correlating metabolic activity and malignancy (40, 41, 48). To further validate measurement of metabolic activity, knockout studies were conducted by inhibiting metabolism within adherent cells using Triton X-100, a cell membrane permeabilizing agent. Cells treated with the detergent demonstrated significantly reduced intracellular dynamics, validating the biological nature of measured intracellular dynamics (Supplemental Fig. S5).

Compared to BCARS or other label-free chemical imaging methods, deep-UV microscopy offers a simper and faster approach for longitudinal imaging of live cells. In this study, we demonstrate widefield imaging up to 100 Hz with sufficient SNR for dynamic analysis, capturing intracelllar dynamics in seconds, offering higher throughput imaging compared to point-scanning techniques. Importantly, the observed dynamics do not appear to rely on a strictly coherent illumination mechanism. While the experiments conducted in this study were performed with a benchtop system featuring a partially coherent laser-driven plasma source, similar analyses may be feasible in more compact UV systems in which partial spatial coherence is introduced using LED-based illumination. Preliminary experiments using simple incoherent LED illumination revealed measureable intracellular dynamics, although the resulting images did not fully reproduce the spatial detail or image quality achieved with the benchtop plasma-source system (Supplemental Fig. S6). These results suggest that future low-cost, portable deep-UV microscopy systems could be developed for longitudinal monitoring of adherent cell cultures, provided that illumination coherence is optimized. Such systems could be used in a variety of settings, including high-throughput cell analysis during cell therapy manufacturing pipelines, or within incubators for longitudinal monitoring of adherent cell cultures. We note that we did find a dependence of measured intracellular dynamics on optical resolution. All of the work presented in this study was acquired using a 0.5 NA objective, corresponding with a lateral resolution of approximately 300 nm. When time series were blurred and downsampled (e.g., by a factor of two to an effective resolution of 600 nm), notable differences were observed in the resulting dynamic images (Supplemental Fig. S7) as compared to the original data (Fig. 3). While the phasor plots did not vary significantly, the worse spatial resolution was reflected in the pseudo-colorized images that lacked the granular detail of subcellular dynamics demonstrated in this work, thus reducing the spatial specificity and phenotyping ability of this technique.

One primary limitation of this study is the requirement for healthy, well-adhered cells to ensure sufficient subcellular contrast in the collected transmission images. While previous studies have demonstrated deep-UV microscopy for imaging smaller suspension cells (e.g., T cells) (62), adherent imaging provides increased contrast in larger epithelial cells and reduced motion artifacts during longitudinal acquisitions. For translation to high-throughput applications that require imaging of large cells in suspension, microfluidic devices or weak adherence substrates can be used to sufficiently flatten cells for analysis. Furthermore, faster acquisition speeds beyond the 100 Hz imaging rate will require subsequently higher deep-UV illumination power, which can induce phototoxic effects, depending on total acquisition times. We have indeed previously quantified UV fluence thresholds (including for RWPE-1 and WPE1-NB26 cells) required for UV-induced photodamage, and demonstrated that longitudinal live cell imaging studies with UV micrscopy are not hampered by UV-induced photodamage given the low illumination powers used (e.g., 10 µW) and appropriate UV dose fractionation (e.g., short and low duty cycle exposure intervals) (33).

In conclusion, we present deep-UV microscopy as a powerful technique capable of capturing biomolecular contrast, subcellular structure, and dynamic, biomolecule-specific intracellular aactivity. The label-free, simple, and high-throughput nature of the technique enables broad implementation of the technology in the analysis of phenotype-specific dynamic activity. In the future, this could include classification of malignancy in other unique cell types or excised tissues, further investigation of intracellular phenotype-specific responses to drugs, and even discovery of new high-speed biological processes. As a versatile, high-resolution imaging tool, deep-UV microscopy has the potential to serve as a powerful research tool for basic and translation studies of cell state and function.

## Materials and Methods

### Cell culture and sample preparation

RWPE-1, WPE1-NB11, and WPE1-NB26 cells were cultured according to ATCC and literature recommended protocols. All cell lines were maintained in Keratinocyte Serum-Free Medium (17005042; ThermoFisher) supplemented with bovine pituitary extract and recombinant human epidermal growth factor. Cells were cultured in T75 tissue culture flasks and incubated at 37°C with 5% CO_2_. Prior to imaging, cells were seeded in poly-D-lysine coated custom petri dishes with quartz-bottom imaging wells (CGQ-0660; Chemglass) at a concentration of ~20,000 cells/cm^2^ and allowed to adhere overnight. For imaging experiments, cells were rinsed with PBS and placed in fresh medium to minimize movement of debris.

### Deep-UV microscopy workflow

For deep-UV microscopy, each petri dish was placed in the optical path of a custom benchtop deep-UV microscopy. This system, extensively described in prior works, consists of a broadband laser-driven plasma light source (EQ-99X; Energetiq), narrowband bandpass filters corresponding with biomolecular absorption peaks in the UV region (Single Bandpass Filters; Chroma), a 40X UV objective (LMU-40X-UVB; Thorlabs), and a UV-sensitive camera (pco.panda; Excelitas). A schematic of the microscope setup is provided in the Supplemental Information (Supplemental Fig. S7). Additional data with an incoherent LED source (PKB-H50-F35; LaserComponents) placed in the optical path was collected for comparison. Image stacks were captured at 1, 10, and 100 Hz frame rates with 500 frames per stack unless otherwise noted. For long-duration acquisitions, 10,000 frame stacks were collected at 100 Hz.

### BCARS and fluorescence imaging and analysis

Broadband coherent anti-Stokes Raman scattering (BCARS) imaging was used to validate intracellular structures observed in UV microscopy. RWPE-1 cells were fixed in 4% paraformaldehyde for 15 minutes after UV microscopy and imaged using a previously demonstrated BCARS system, equipped with a brightfield microscope to find corresponding field-of-views (12– 14). The data processing pipeline for Raman signal retrieval from BCARS spectra, including singular value decomposition, Kramers-Kronig transfer, phase retrieval, and baseline correction, are detailed in these works as well. Point spectra were then extracted from specific subcellular regions of interest for biochemical validation.

For fluorescent staining of mitochondria and nuclei, a two-step protocol was performed for live-cell labeling with a mitochondria staining, followed by fixation, permeabilization, and nuclear staining with DAPI. MitoTracker Red CMXRos (Thermofisher; M7512) was reconstituted in anhydrous DMSO to a 1 mM stock concentration and diluted to a 200 nM working solution in Keratinocyte SFM. A 1 mL aliquot of MitoTracker solution was added to each coverslip and incubated at 37°C for 30 minutes to allow mitochondrial uptake of the dye. Following incubation, cells rinsed twice with pre-warmed PBS and then fixed with 3.7% formaldehyde in PBS for 15 minutes at 37°C. After fixation, cells were washed three times with PBS and subsequently permeabilized with 0.2% Triton X-100 (FisherScientific; BP 151-100). For nuclear labeling, cells were incubated with 1 µg/mL DAPI (Sigma Aldrich; MBD0015) for 10 minutes at room temperature, protected from light. Cells were then washed two times with PBS and stored in PBS at 4C until imaging. Imaging was performed with a confocal laser scanning microscope (LSM 900; Zeiss) with a 63X 1.4NA oil immersion objective.

### UV image analysis and feature analysis

Raw UV transmission images were converted to attenuation images by computing the negative logarithm of background-corrected intensity values. Two complementary dynamic analyses were performed. First, the time series for each pixel within segmented cells was extracted. Pixelwise power spectral density (PSD) plots were computed via the frequency response of each time series, and then the log-log slope of each PSD was fit to quantify frequency-dependent dynamic activity. Extremely low frequency (e.g., <0.1 Hz) activity was excluded as a linear model was not followed in this regime, such that the frequency range for fitting the log-log slope of each PSD was approximately 0.1 Hz to half the imaging frequency. Phasor analysis was also performed during which the frequency response is decomposed into two terms, denoted g and s, representing the real and imaginary parts of the signal’s Fourier Transform, respectively, which also serve as quantified metrics for pixelwise fluctuation. Pseudo-colorized images were then created using the extracted pixelwise dynamic values with blue and red representing low and high activity, respectively. In short, HSV maps were developed using pixelwise dynamics as the hue channel (i.e., phasor coordinate or fitted power law log slope value), a matrix of ones as the saturation channel, and UV attenuation as the value channel, enabling visualization of both structure and measured dynamics. The upper bounds of the dynamic activity ranges (e.g., 0.5-5 Hz for 10 Hz imaging rate and 5-50 Hz for 100 Hz imaging rate) were set using the Nyquist limit (1/2 of the sampling rate), while the lower bounds were set by requiring at least approximately 30 cycles within each time series for minimum resolvable frequency analysis, shown to be sufficient in recent literature (11, 63) for quantifying cellular dynamics.

Quantitative features were extracted from both static and dynamic images. Static features included morphological properties (e.g., area, circularity) and textural metrics (e.g., entropy, homogeneity) computed from UV attenuation images at a given wavelength. Dynamic features included mean cellwise phasor g and s values, fitted power law slope coefficients, and texture features calculated from dynamic feature maps. These features have been extensively detailed in previous works for tissue and cell characterization, and in total over 1,000 features were calculated for each cell.

To determine significant features for phenotyping, we performed chi-squared feature ranking on the extracted feature set. Eight features were used for subsequent visualization and classification, comprising both static and dynamic features. Statistical comparisons among cell types were performed using two-tailed unpaired *t*-tests. Each bar plot shown in the manuscript includes error bars corresponding with standard deviation. Uniform Manifold Approximation and Projection (UMAP) was also used to visualize separation among the three cell lines with typical parameters (e.g., nearest neighbors ~15, minimum distance between points ~0.1).

## Acknowledgements

We gratefully acknowledge the following funding sources: National Institutes of Health National Institute of General Medical Sciences: R35GM147437 (F.E.R); Burroughs Welcome Fund grant: CASI BWF 1014540 (F.E.R.); National Science Foundation grant: NSF CBET CAREER 1752011 (F.E.R.); National Institutes of Health National Institute of Biomedical Imaging and Bio-engineering grant: R41-EB035057 (F.E.R.); National Institutes of Health National Heart, Lung, and Blood Institute grant: R43-HL167435 (F.E.R.); Georgia Institute of Technology;

## Author contributions

V.G., N.T., M.C., and F.E.R. designed the experiments outlined in this work. V.G. and M.S. prepared samples for imaging. V.G., M.S., and N.T. acquired data. V.G. and N.T. prepared and performed data analysis. V.G., N.T., and F.E.R prepared all figures and the manuscript. All authors reviewed and approved the final version of the manuscript. Fig. 1 created with BioRender.com.

## Data availability

Data underlying the results presented in this paper are not publicly available at this time but may be obtained from the authors upon reasonable request.

