## Supplemental Information for "Deep-ultraviolet microscopy reveals biomolecular spatiotemporal intracellular dynamics"

bioRxiv

### Supplemental information for: Deep-ultraviolet microscopy reveals biomolecular specific spatiotemporally variant intracellular dynamics

**Viswanath Gorti,[a] Mingxuan Si,[a] Nathan Taylor,[b] Marcus Cicerone,[b] and Francisco E. Robles[a,*]**
[a]Wallace H. Coulter Department of Biomedical Engineering, Georgia Institute of Technology and Emory University, Atlanta, Georgia, USA
[b]Department of Chemistry and Biochemistry, Georgia Institute of Technology, 950 Atlantic Drive, Atlanta, Georgia 30332, USA
[*]

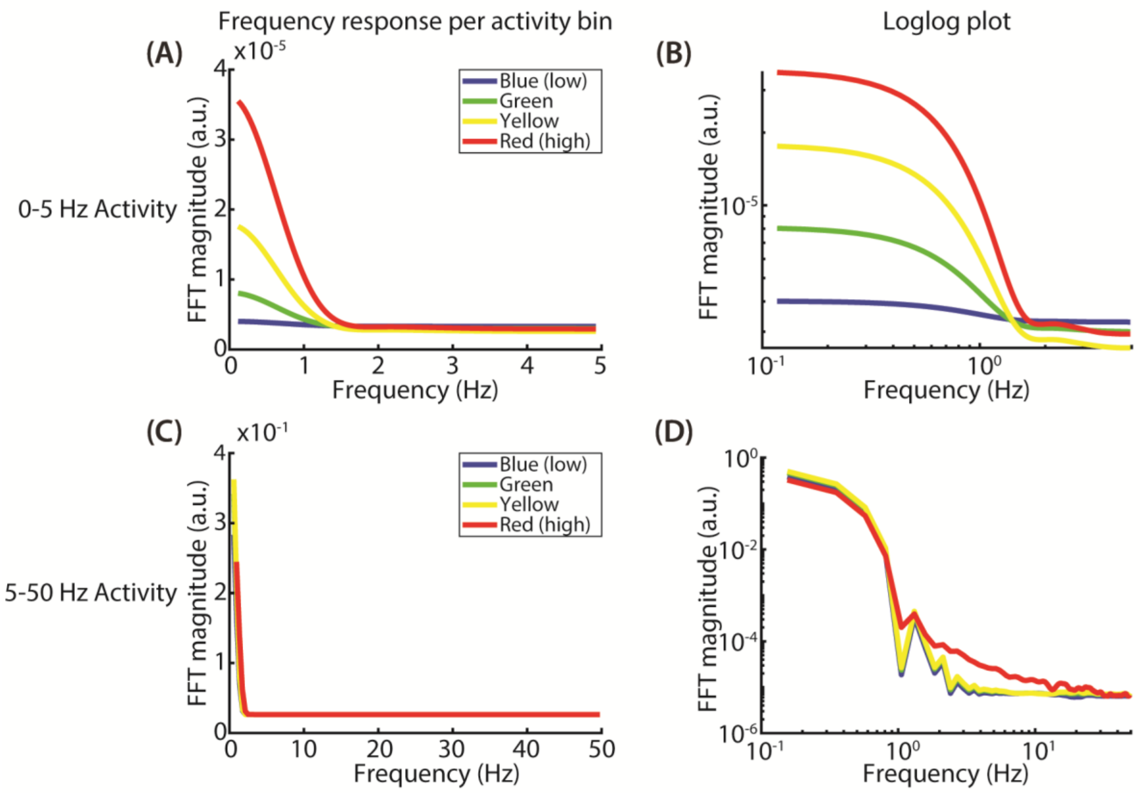


**Fig. S1. Frequency-domain dynamics grouped by activity level and imaging frame rate.** (A-B) Frequency responses of RWPE-1 cells imaged at 10 Hz using 255 nm illumination, separated into four activity bins (blue = low, green = medium-low, yellow = medium-high, red = high) based on their phasor analysis quantified activity. In this analysis, the activity bins correspond with quartiles of quantified activity. (A) Linear-scale frequency response for the 0-5 Hz activity range showing increasing low-frequency power with higher activity levels, consistent with stronger slow intracellular fluctuations. (B) Corresponding log-log spectra showing relative spectral decay rates across activity bins. (C-D) Frequency responses of RWPE-1 cells imaged at 100 Hz using 255 nm illumination for the 5-50 Hz activity range. (C) Linear-scale frequency response for the 5-50 Hz activity range showing rapid drop-off in high-frequency power across all activity bins. (D) Log-log spectra revealing increased high-frequency activity for pixels in the highest activity (red) bin.

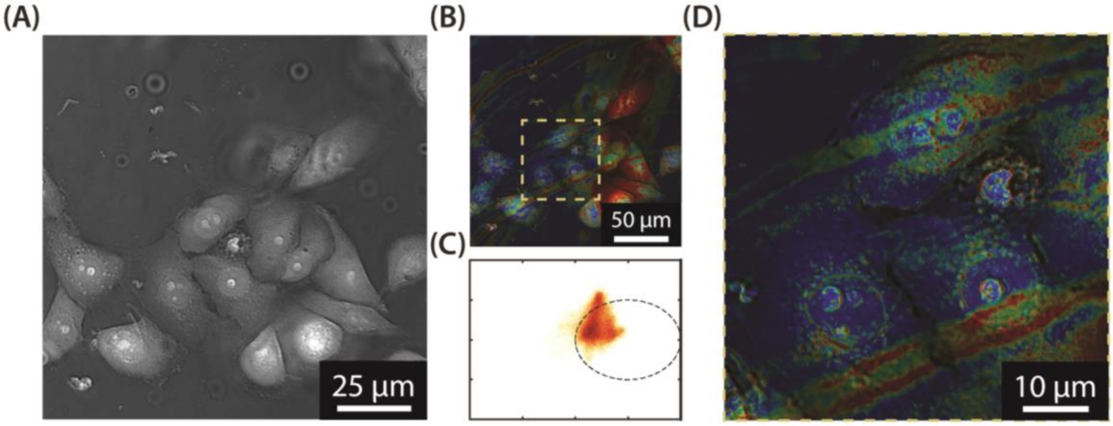


**Fig. S2. Limitations of imaging at low (1 Hz) frame rate.** (A) UV attenuation image at 239 nm illumination of live RWPE-1 cells. (B) Phasor-based pseudo-colorized map computed using a 1 Hz 500 frame time series (inset in D) showing confounding signal from out of focus debris floating through the field of view during acquisition. (C) Phasor plot from the field of view in (B) demonstrating noisy distribution of phasor values different from typical intracellular signal.

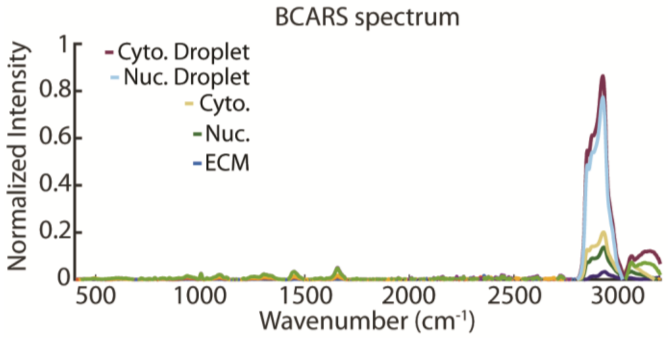


**Fig. S3. Full BCARS spectra for significant intracellular regions.** BCARS spectra shown from 400 – 3200 $cm^{-1}$ for pixels corresponding with the extracellular matrix (ECM), cell nucleus, cytoplasm, a nuclear lipid droplet, and a cytoplasmic lipid droplet.

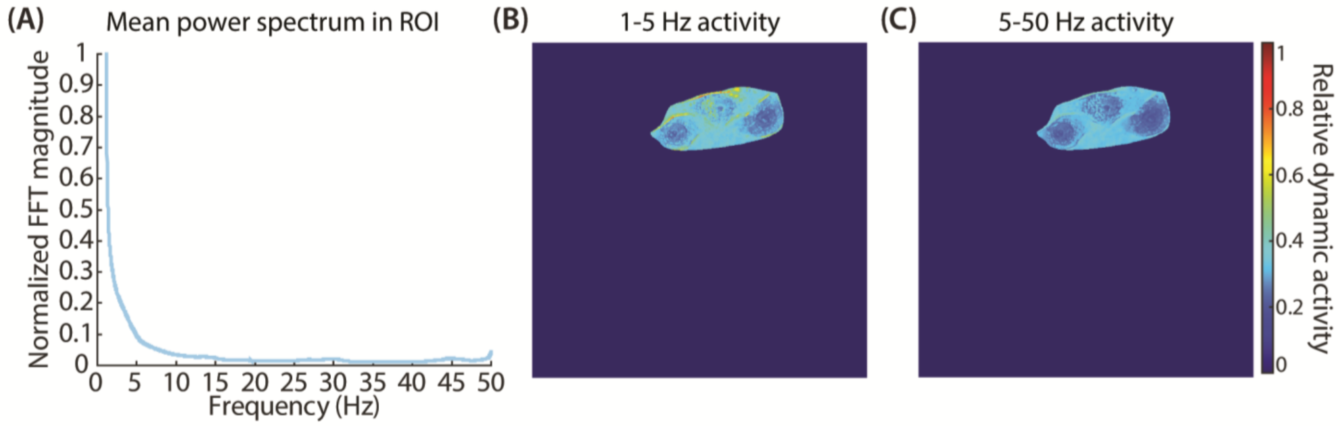


**Fig. S4: Extended duration deep-UV time series captured at 239 nm.** (A) Normalized mean power spectrum averaged for all pixels within three RWPE-1 cells, showing decaying intensity with no dominant frequency components. (C,D) Spatial maps of pixelwise dynamic activity in the selected region for two frequency ranges: 1-5 Hz (C) and 5-50 Hz (D). Activity was quantified by averaging the signal magnitude within each frequency range and then plotting values to their corresponding spatial locations.

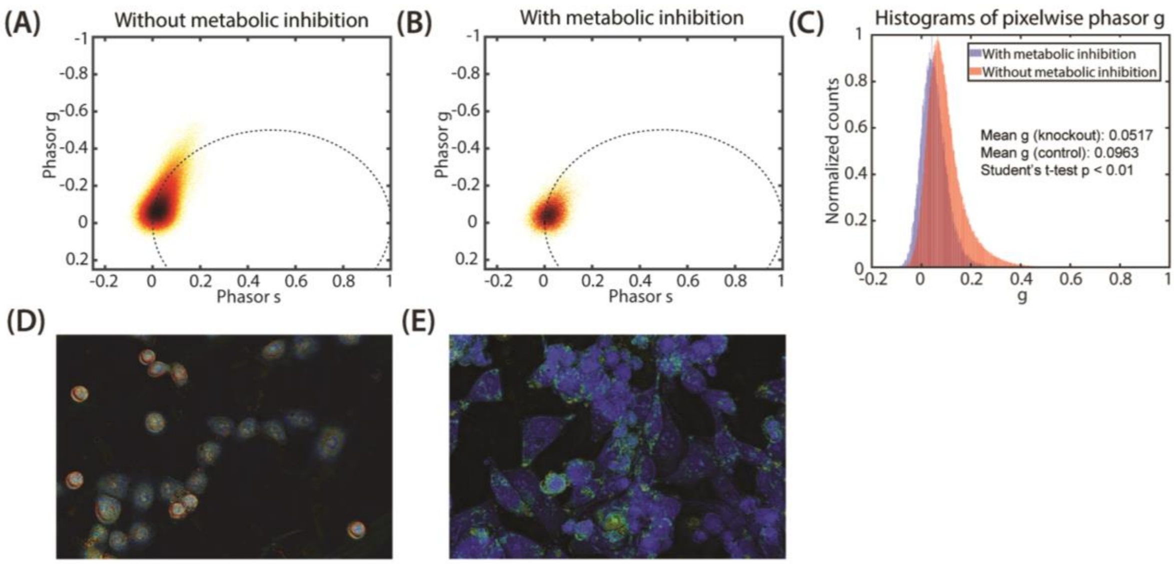


**Fig. S5. Metabolic inhibition of intracellular dynamics reduces measured signal.** (A-B) Phasor plots of RWPE-1 cells imaged at 10 Hz with 255 nm illumination without (A) and with (B) metabolic inhibition using 0.05% Triton X-100 added to the imaging medium 5 minutes prior to imaging. (C) Histogram of pixelwise phasor g values showing a decrease in mean g value for cells treated with the inhibitor. (D-E) Phasor-colorized images of cells without (D) and with (E) metabolic inhibition, showing a decrease in measured dynamic activity.

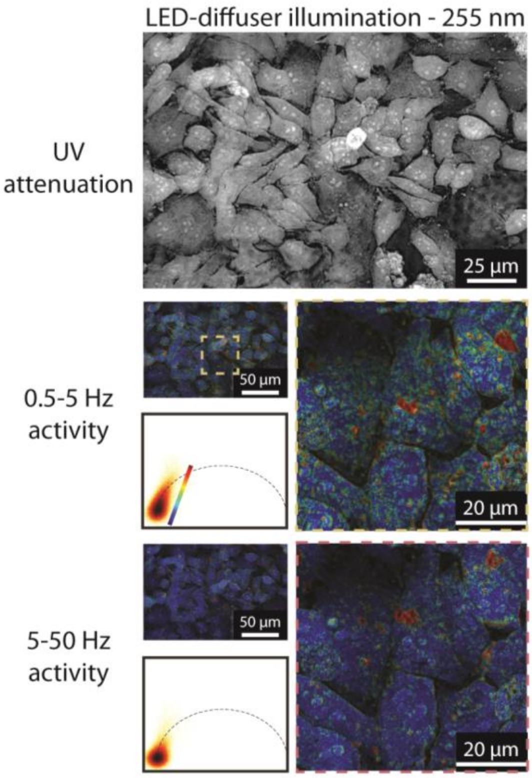


**Fig. S6. Dynamics of RWPE-1 cells from deep-UV image stacks at incoherent (LED) 255 nm illumination.** UV attenuation image (top) and phasor pseudo-colorized images shown at low (middle) and high (bottom) frequency imaging.

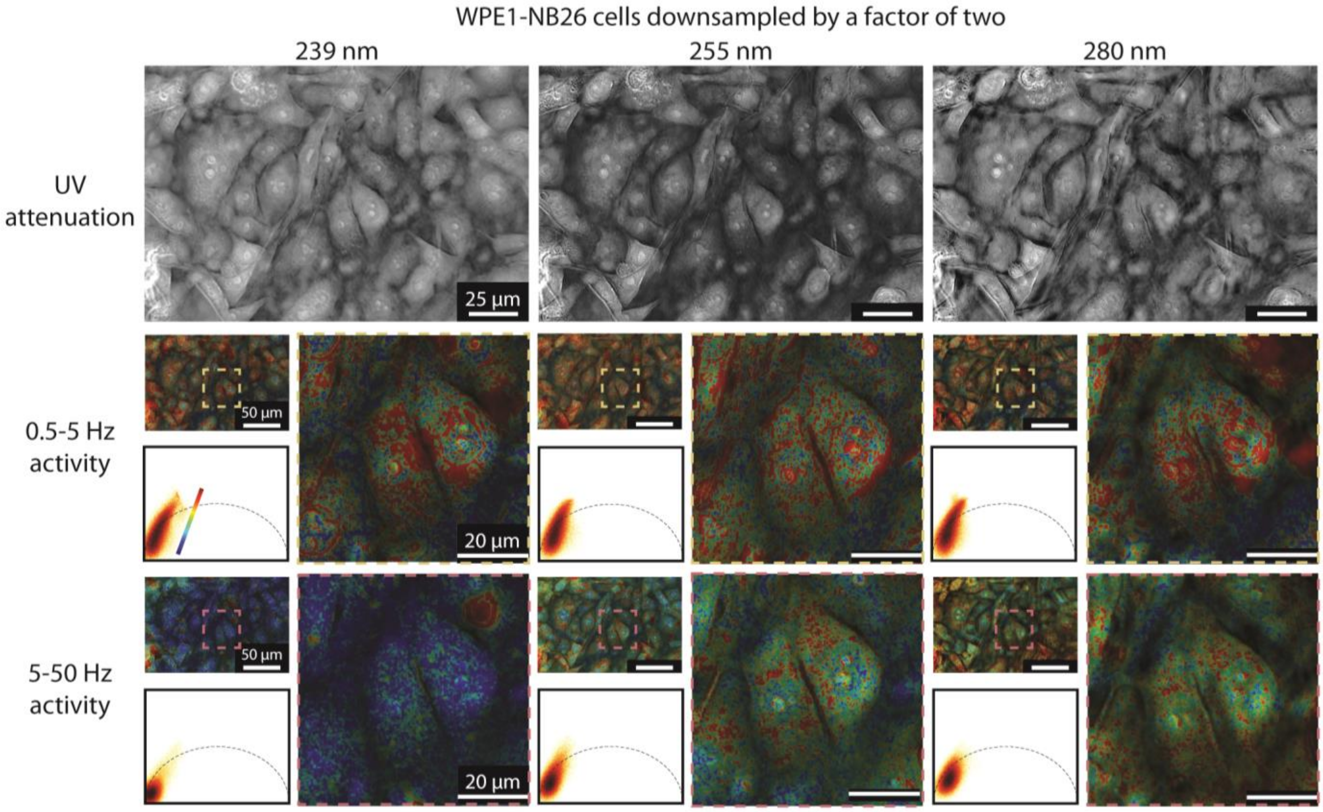


**Fig. S7. Effect of spatial resolution on measured dynamic intracellular activity.** WPE1-NB26 cell data shown in Fig. 3 after Gaussian blurring spatial downsampling of the original image stacks by a factor of two. Data includes UV attenuation images of the field of view at each illumination wavelength (top row), along with phasor plots and pseudo colorized images corresponding with 0.5-5 Hz (middle row) and 5-50 Hz (bottom row) activity. Compared to Fig. 3, the reduced resolution results in lower spatial specificity and increased noise through the cells.

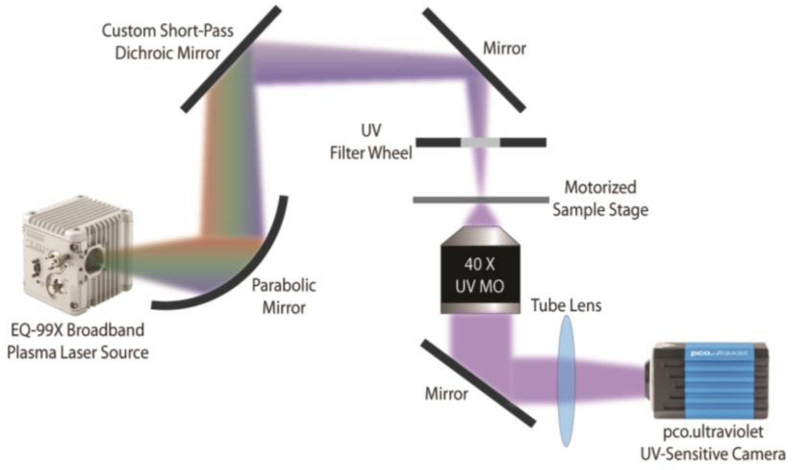


**Fig. S7. Benchtop deep-UV microscopy system used in this work.** Schematic of our previously demonstrated deep-UV microscopy, comprising a broadband plasma laser source, narrowband UV band-pass filters, a UV objective, and scientific UV-sensitive camera.
